# JWA deficiency accelerates aging through disrupting intestinal epithelial homeostasis via Notch1/PPARγ/Stat5 axis

**DOI:** 10.1101/2022.01.17.476552

**Authors:** Xiong Li, Jingwen Liu, Luman Wang, Yan Zhou, Yifan Wen, Kun Ding, Lu Zou, Xia Liu, Aiping Li, Yun Wang, Heling Fu, Min Huang, Guoxian Ding, Jianwei Zhou

**Affiliations:** Department of Molecular Cell Biology & Toxicology, Center for Global Health, School of Public Health, Nanjing Medical University, Nanjing 211166, China; Key Laboratory of Modern Toxicology of Ministry of Education, School of Public Health, Nanjing Medical University, Nanjing, 211166, China; Jiangsu Key Lab of Cancer Biomarkers, Prevention and Treatment, Collaborative Innovation Center for Cancer Medicine, Nanjing Medical University, Nanjing 211166, China; Animal Core Facility of Nanjing Medical University, Jiangsu Animal Experimental Center of Medical and Pharmaceutical Research, Nanjing 211166, China; Department of Geriatrics, Division of Geriatric Endocrinology, The First Affiliated Hospital of Nanjing Medical University, Nanjing 210029, China

**Keywords:** JWA, Aging, Intestinal regeneration, Intestinal stem cell, Notch signal

## Abstract

Aging usually suppresses the renewal and regeneration of intestinal epithelium. The imbalance of intestinal epithelial homeostasis may also be a promoter for aging. JWA responds to oxidative stress and repairs damaged DNA; it participates in multiple cellular processes like cell proliferation and differentiation. Here we identified JWA as a new aging-associated gene, whose deletion-accelerated aging in mice was related to intestinal epithelium atrophy. We further knocked out intestinal epithelial JWA and found it disrupted intestinal epithelial homeostasis, thus promoting aging in mice. Mechanistically, we discovered that JWA deficiency promoted Notch1 ubiquitination degradation via ERK/Fbxw7 cascade and interfered with the PPARγ/Stat5 signal axis. This reduced the intestinal stem cell function and altered the intestinal epithelial cell lineage distribution, finally suppressing the renewal and regeneration of intestinal epithelium. Our results demonstrated that JWA is a new aging-associated gene essential for the renewal and regeneration of intestinal epithelium. We also provide a new idea that maintaining intestinal epithelial homeostasis may be a potential anti-aging strategy in humans or mammals.

## Introduction

The precise regulation of tissue renewal and regeneration is an effective mechanism to prevent and slow down tissue degeneration (Picerno *et al*, 2021). In mammals, the intestinal epithelium renews rapidly and usually turns over within 3-5 days, due to the organized cell proliferation, differentiation, migration, and apoptosis, as well as the regenerative capability of intestinal stem cells (ISCs) (Xiao *et al*, 2019). Unfortunately, these characteristics make intestinal epithelium more susceptible to external environmental factors like ionizing radiation. The high dose radiation exposure can induce intestinal epithelial injury followed by gastrointestinal syndrome (GIS). The ISCs respond immediately and regenerate quickly to repair the injured intestinal epithelium (Chaves-Perez *et al*, 2019). Therefore, certain factors that negatively affect the ISC maintenance can inevitably inhibit renewal and prevent regeneration of intestinal epithelium after injury.

Aging means the age-dependent, progressive and irreversible decline in physiological functions that eventually leads to death. The degradation of various tissues and organs usually accompanies the aging process, making aging one of the main risk factors for human pathologies such as cancer, diabetes, cardiovascular disorders, and neurodegenerative diseases (Lopez-Otin *et al*, 2013). The turnover of intestinal epithelium often decreases with age due to the functional declines of ISC and supporting niche cells, causing slower renewal and regeneration in aging intestinal epithelium after irradiation-induced damage (Pentinmikko *et al*, 2019).

On the other hand, intestinal epithelium plays an essential role in nutrient absorption; it is also the primary immune barrier to prevent lumen microorganisms and toxic substances from entering the circulation. The imbalance of nutrient intake through the intestine epithelium can promote aging and affect health of the elderly (Sun *et al*, 2021), although the dietary restriction (DR) without malnutrition has many profound beneficial effects on longevity (Green *et al*, 2022; Wang *et al*, 2022). Meanwhile, aging is also potentially related to intestinal microbial imbalance and inflammation (Shrout *et al*, 2021). Therefore, loss of intestinal epithelial integrity may be the driver for aging and obstacle to healthy aging. Intestinal epithelial homeostasis alteration can also influence the lifespan of lower creatures such as nematode and *Drosophila* (Sasaki *et al*, 2021; Yang *et al*, 2021), making the intestine considered a potential target organ for anti-aging (Rera *et al*, 2013). However, limited evidence proves that intestinal epithelial disruption accelerates aging, especially in mammals.

JWA, also named ADP-ribosylation-like factor 6 interacting protein 5 (Arl6ip5), is initially cloned from human bronchial epithelial (HBE) cells induced by all-trans retinoic acid. We previously demonstrated that JWA responded to oxidative stress and participated in DNA single-strand damage repair (Chen *et al*, 2007b; Wang *et al*, 2009), the unrepaired DNA damage accumulation-induced genome instability is one of the hallmarks of aging (Lopez-Otin et al., 2013). We further found that JWA exerted neuroprotective role in the mouse model of Parkinson’s disease via alleviating oxidative stress and inhibiting inflammation (Wang *et al*, 2018; Zhao *et al*, 2017). However, there is not enough exploration and evidence to prove whether JWA is involved in aging. Moreover, JWA participates in multiple cellular processes like cell proliferation, differentiation, and apoptosis (Chen *et al*, 2004; Li *et al*, 2003), making it capable of regulating intestinal epithelial renewal and regeneration. Therefore, it is an exciting hypothesis that JWA regulates intestinal epithelial renewal and regeneration, thus affecting the aging process.

In this study, we demonstrated that JWA was essential for the renewal and regeneration of intestinal epithelium, and for the first time we claimed that JWA deficiency accelerated aging through disrupting intestinal epithelial homeostasis in mice. Our results will also suggest the maintenance of intestinal epithelial homeostasis as a potential anti-aging strategy.

## Results

### JWA is a new aging-associated gene

To confirm the relationship between JWA and aging, we firstly examined JWA in the spleens and livers of mice and found the old mice had lower JWA levels than young mice (Fig. 1A, B, Appendix Fig. S1A and B). We previously generated the JWA knockout (JWA^ko^) mice, and in the present study we observed that JWA^ko^ mice exhibited thinner bodies (Fig. 1C) and lower body weight than wild type (JWA^wt^) mice starting from 4-week-old (Fig. 1D). Subsequently, we built a mice cohort for survival analysis, the survival curves depicted that the medium lifespan of JWA^ko^ mice was only 290.0 days, which was significantly shorter than JWA^wt^ mice (Fig. 1E). We performed a series of phenotypic analyses in mice at the young and relatively old stage and found multiple aging-related phenotypes in both young and old JWA^ko^ mice, such as lost hair (Appendix Fig. S2A and S3A), deformed spine (Appendix Fig. S2B, C, S3B and C), disordered cardiovascular function (Appendix Fig S2D-G and S3D-G), and reduced basal metabolism (Appendix Fig. S2H-K and S3H-K). Furthermore, we observed elevated senescence-associated β-galactosidase (SA-β-gal) positive cells in the JWA^ko^ mice liver sections (Fig. 1F), indicating the cellular senescence. Our results suggested that JWA was a new aging-associated gene whose deletion accelerated mice aging.

**Figure 1.**
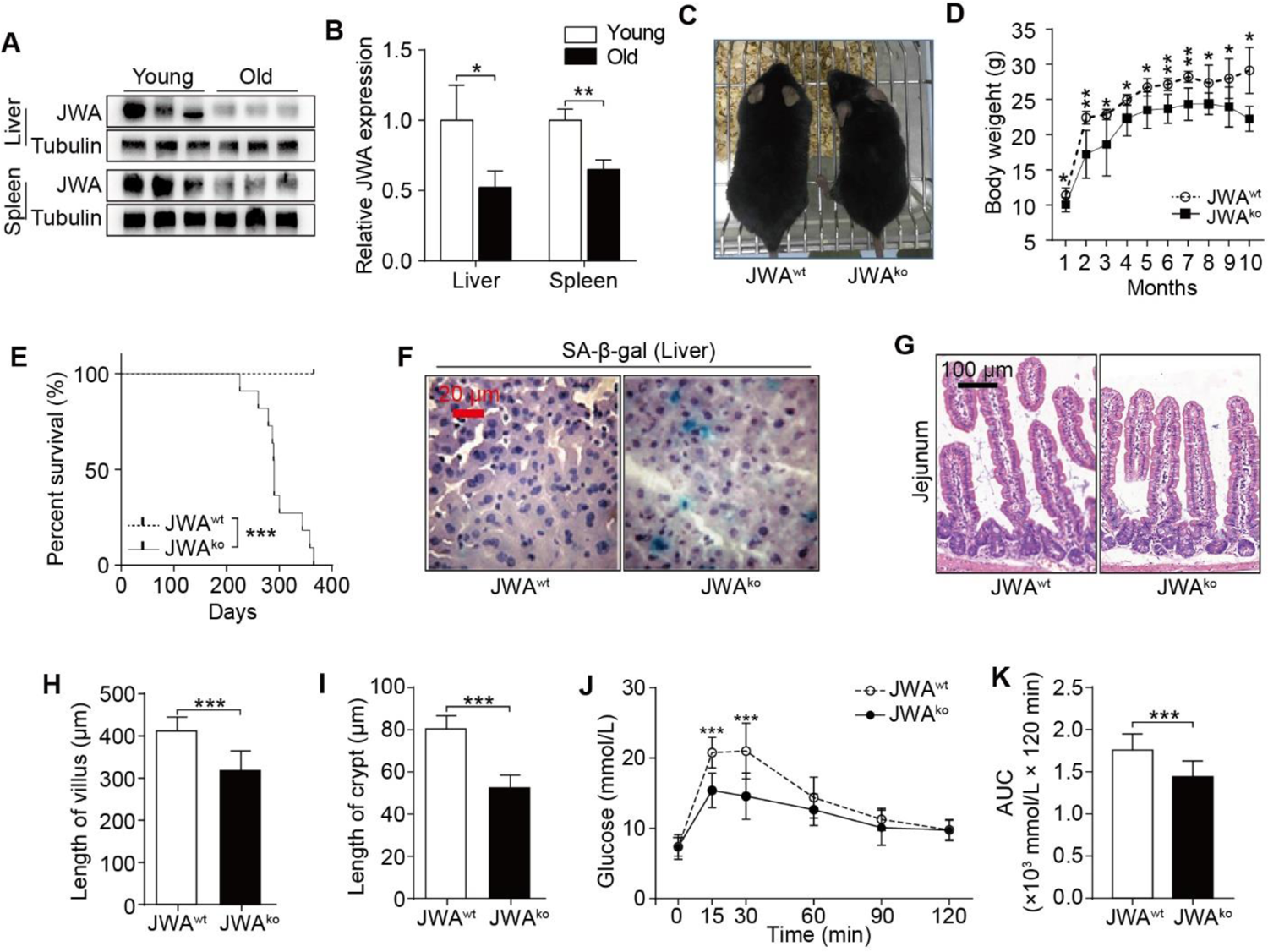
JWA is a new aging-associated gene whose deletion accelerates aging through disrupting intestinal epithelial homeostasis. A, B. Immunoblotting of JWA (A) and relatively JWA levels (B) in the liver and spleen of young (2-month-old, n=3) and old (20-month-old, n=3) mice. C. Representative photograph of 6-month-old JWA^wt^ and JWA^ko^ mice. D. Body weight curve of JWA^wt^ and JWA^ko^ mice from 1-month-old to 10-month-old; n=6 for each genotype. E. Survival curve of JWA^wt^ and JWA^ko^ mice; n=12 for JWA^wt^ mice and n=11 for JWA^ko^ mice. F. SA-β-gal staining in the liver sections of JWA^wt^ and JWA^ko^ mice. Scale bar: 20 μm. G-I. H&E staining of the jejunum sections (K), and the length measurement of villi (L) and crypts (M) in 6-month-old JWA^wt^ and JWA^ko^ mice; n=3 for each genotype; scale bar: 100 μm. J, K. OGTT curve (N) and the AUC measurement (O) in 6-month-old JWA^wt^ and JWA^ko^ mice; n=12 for each genotype. Data information: In (B, D, H-K), data are presented as mean ± SD, \**P* <0.05, \*\**P* <0.01 and \*\*\**P* <0.001 (Student’s *t*-test). In (E), \*\*\**P* <0.001 (Log-Rank test).

### JWA deletion-accelerated aging originates from the intestine epithelium

We explored the histomorphology of the principal organs of mice (Fig. 1G and Appendix Fig. S4) and found that the primary alteration was the atrophy of intestinal epithelium (Fig. 1G), which showed the reduced length of villi and crypts (Fig. 1H and I). We evaluated the intestine epithelium absorption through oral glucose tolerance test (OGTT) and found that JWA^ko^ mice exhibited poor glucose absorption (Fig. 1J and K). Moreover, JWA^ko^ mice consumed less food and water than JWA^wt^ mice (Appendix Fig. S2H, I, S3H and I). These results further reminded us of the probable alterations of intestinal epithelial structure and function due to the JWA deficiency. Furthermore, we observed that JWA deletion delayed the development of intestinal epithelium in both embryonic (Fig. EV1A-D) and newborn mice (Fig. EV1E). However, there was no difference in the embryonic weight between JWA^ko^ and JWA^wt^ mice (Fig. EV1F). These results alluded that the intestine epithelium might be the originate of JWA deletion-accelerated aging.

### JWA expresses higher in intestinal crypts than villi

Through The Human Protein Atlas database (https://www.proteinatlas.org), we got that the JWA mRNA and protein expressed widely in the human tissues and organs (Appendix Fig. S5A and B). JWA protein was highest existing in the human duodenum and the whole small intestine (Appendix Fig. S5B and C). We also detected that the JWA protein exhibited high levels in mouse intestine (Appendix Fig. S5D), especially the middle intestine (jejunum) (Appendix Fig. S5E and F). Moreover, we observed that the protein level of JWA was higher in intestinal crypts than villi (Fig. 2A). Therefore, we hypothesized that JWA might play a critical role in renewal and regeneration of intestinal epithelium, owing to the intestinal stem cells (ISCs) and niche cells located in the crypts, which are involved in the maintenance of intestinal epithelium.

**Figure 2.**
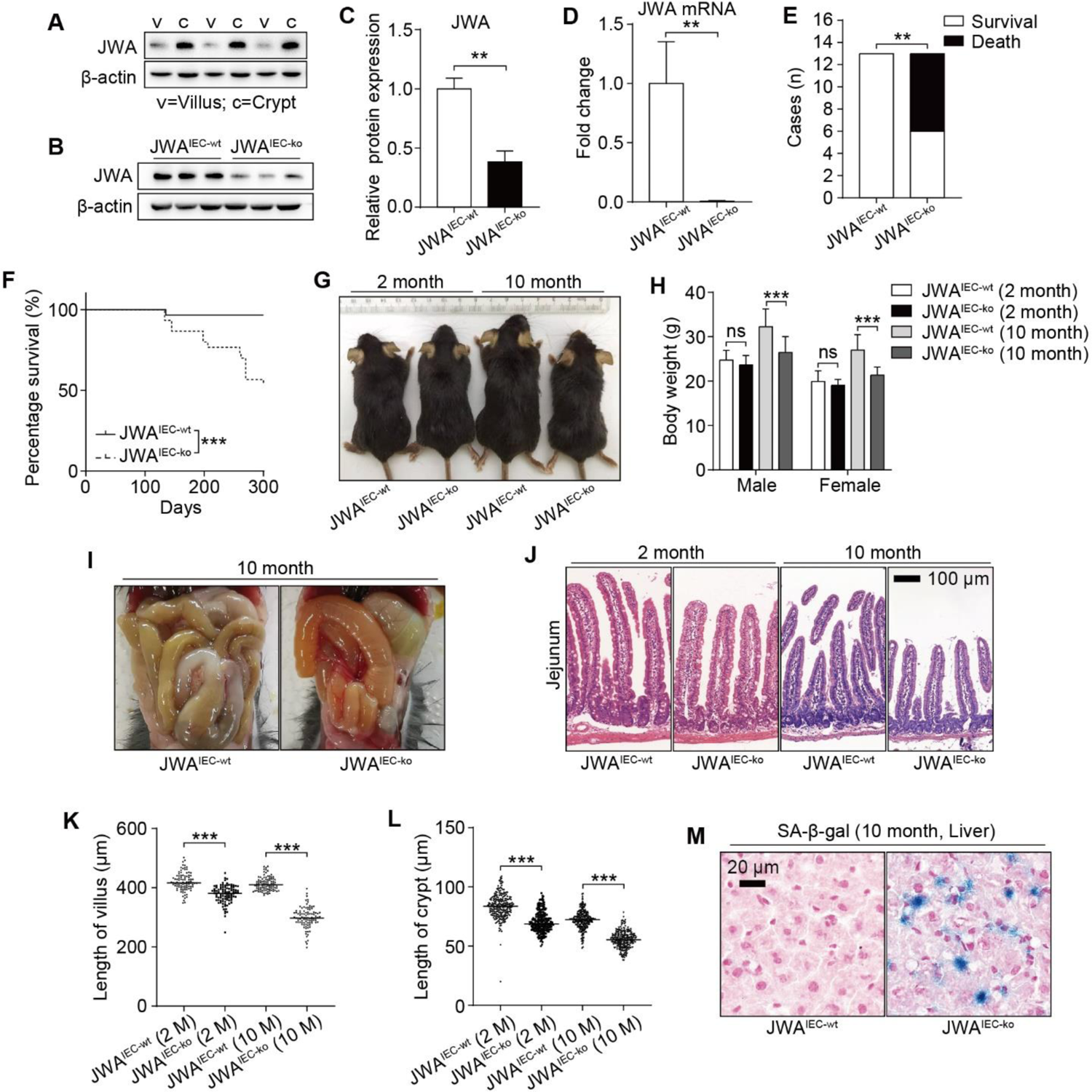
Intestinal epithelial JWA deletion disrupts intestinal epithelial homeostasis and accelerates mice aging. A. Immunoblotting of JWA in intestinal villi and crypts of wild-type mice; n=3, “v” represents “villi”; “c” represents “crypts”. B, C. Immunoblotting of JWA protein (B) and relatively JWA levels (C) in crypts of JWA^IEC-wt^ and JWA^IEC-ko^ mice; n=3 for each genotype. D. QRT–PCR detection of JWA levels in crypts of JWA^IEC-wt^ and JWA^IEC-ko^ mice; n=3 for each genotype. E. Statistics of death and survival JWA^IEC-wt^ and JWA^IEC-ko^ mice by the end of 10-month-old; n=13 for each genotype at initial. F. Survival curve of JWA^IEC-wt^ and JWA^IEC-ko^ mice by the end of 10-month-old; n=30 for each genotype. G. Representative photograph of JWA^IEC-wt^ and JWA^IEC-ko^ mice at 2-month-old and 10-month-old respectively. H. Body weight of JWA^IEC-wt^ and JWA^IEC-ko^ mice at 2-month-old and 10-month-old respectively; n=6-11 for each genotype. I. Representative abdominal cavity images of JWA^IEC-wt^ and JWA^IEC-ko^ mice at 10-month-old. J-L. H&E staining of the jejunum sections (J), and the length measurement of villi (K) and crypts (L) in JWA^IEC-wt^ and JWA^IEC-ko^ mice at 2-month-old and 10-month-old respectively; n=3 for each genotype; scale bar: 100μm. M. SA-β-gal staining in the liver sections of JWA^IEC-wt^ and JWA^IEC-ko^ mice at 10-month-old. Scale bar: 20 μm. Data information: In (C, D, H, K, L), data are presented as mean ± SD, ^ns^ No significance, \*\**P* <0.01 and \*\*\**P* <0.001 (Student’s *t*-test). In (E), \*\**P* <0.01 (Fisher’s exact test). In (F), \*\*\**P* <0.001 (Log-Rank test).

### Intestinal epithelial JWA deletion destroys the homeostasis of the intestinal epithelium and promotes mice aging

To invest the role of JWA on intestinal epithelial homeostasis maintenance, we constructed the intestinal epithelial JWA knockout (JWA^IEC-ko^) and littermate wild type (JWA^IEC-wt^) mice (Fig. EV2A and B). We confirmed that both the mRNA and protein levels of JWA were significantly down-regulated in crypts (Fig. 2B-D, Fig. EV2C and D). In the mice cohort, we found that most of the JWA^IEC-ko^ mice died before 10-month-old (Fig. 2E), and the survival rate of JWA^IEC-ko^ mice was significantly lower than JWA^IEC-wt^ mice (Fig. 2F). Like JWA^ko^ mice, JWA^IEC-ko^ mice also showed thinner bodies and lighter weights than JWA^IEC-wt^ mice. Interestingly, this only manifested at 10-month-old, rather than 2-month-old (Fig. 2G and H). We dissected the euthanized 10-month-old mice and found that most JWA^IEC-ko^ mice showed intestinal congestion and bloat signs (Fig. 2I). We then observed that JWA^IEC-ko^ mice had shorted villi and crypts than JWA^IEC-wt^ mice, especially at 10-month-old (Fig. 2J-L). Furthermore, to verify our hypothesis that JWA deletion-accelerated aging originated from intestine epithelium, we stained the SA-β-gal positive cells in liver sections from 10-month-old mice and found more senescent cells in JWA^IEC-ko^ mice than JWA^IEC-wt^ mice (Fig. 2M). Moreover, we tested the activity of several redox-related enzymes such as superoxide dismutase (SOD), glutathione peroxidase (GSH-px), and catalase (CAT) in the serum of 10-month-old mice. We found the activities of the three enzymes were all reduced due to intestinal epithelial JWA deletion (Fig. EV2E-G), revealing the deterioration of antioxidant capacity potential, which was one of the hallmarks of aging. Our results suggested that intestinal epithelial JWA deletion could disrupt intestinal epithelial homeostasis, which might be an accelerator for the aging process in mice.

### Intestinal epithelial JWA deletion reduces intestinal stem cells

Intestinal epithelial integrity depends on the proliferation, differentiation, migration of intestinal epithelial cells and function of intestinal stem cells. The ki67 staining showed intestinal epithelial JWA deletion inhibited the proliferation of crypt cells (Fig. 3A, B and Fig. EV3A). It also hindered the intestinal epithelium renewal, revealed by BrdU^+^ cell migration assay (Fig. 2C and D). We next found that intestinal epithelial JWA deletion markedly reduced the ISC number at the intestinal crypts, revealed by the decrease of olfactomedin 4 (olfm4) positive cells (Fig. 3E, F and Fig. EV3B-D). Moreover, reduced mRNA levels of ISC markers such as Leucine-rich repeat containing G protein-coupled receptor 5 (lgr5), olfm4, SPARC-related modular calcium-binding protein 2 (smoc2), Achaete-scute family BHLH transcription factor 2 (ascl2), and Musashi RNA binding protein 1 (msi1) were detected in crypts from JWA^IEC-ko^ mice (Fig. 3G). These results further suggested that JWA deficiency might suppress the renewal and regeneration of intestinal epithelium.

**Figure 3.**
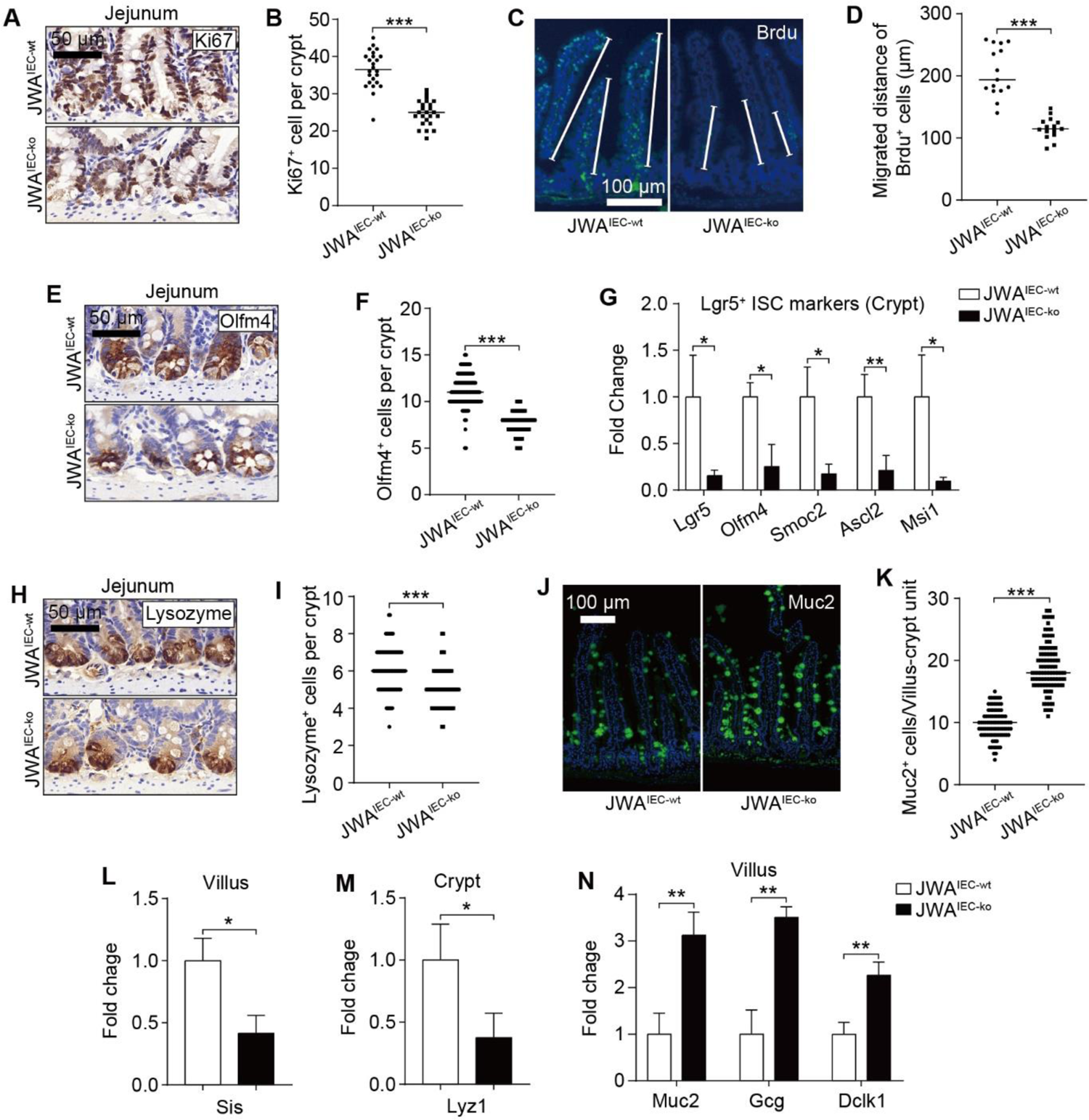
Intestinal epithelial JWA deletion reduces intestinal stem cells and skews the distributions of intestinal epithelial cells lineage. A, B. Immunochemistry staining of ki67 in the intestinal sections of JWA^IEC-wt^ and JWA^IEC-ko^ mice (A), and the ki67 positive cell counts in crypts (B); n=3 for each genotype; scale bar: 50 μm. C, D. Immunofluorescence staining of BrdU in the intestinal sections of JWA^IEC-wt^ and JWA^IEC-ko^ mice (C), and the BrdU positive cells migration measurement (D); n=3 for each genotype; scale bar: 100 μm. E, F. Immunochemistry staining of olfm4 in the intestinal sections of JWA^IEC-wt^ and JWA^IEC-ko^ mice (E), and the olfm4 positive cell counts in crypts (F); n=3 for each genotype; scale bar: 50 μm. G. QRT–PCR detection of ISC markers in the intestinal sections of JWA^IEC-wt^ and JWA^IEC-ko^ mice; n=3 for each genotype. H, I. Immunochemistry staining of lysozyme in the intestinal sections of JWA^IEC-wt^ and JWA^IEC-ko^ mice (H), and the lysozyme positive cell counts in crypts (I); n=3 for each genotype; scale bar: 50 μm. J, K. Immunofluorescence staining of muc2 in the intestinal sections of JWA^IEC-wt^ and JWA^IEC-ko^ mice (H), and the muc2 positive cell counts in villus-crypt units (I); n=3 for each genotype; scale bar: 100 μm. L-N. QRT–PCR detection of the absorption enterocytes marker sis in villi (L), Paneth cell marker lyz1 in crypts (M), goblet cell marker muc2, enteroendocrine cell marker gcg, and tuft cell marker dclk1 in villi (N) of JWA^IEC-wt^ and JWA^IEC-ko^ mice; n=3 for each genotype. Data information: In (B, D, F, G, I, K-N), data are presented as mean ± SD, \**P* <0.05, \*\**P* <0.01 and \*\*\**P* <0.001 (Student’s *t*-test).

### Intestinal epithelial JWA deletion skews the distributions of intestinal epithelial cells lineage

The distribution balance of IEC lineages also plays an essential role in intestinal epithelial homeostasis maintenance. To explore the impact of JWA deficiency on IEC lineages, we examined the levels of various IEC markers. Results showed markedly decreased lysozyme positive Paneth cells number in crypts (Fig. 3H, I and Fig. EV3G), elevated muc2 and Alcian blue positive goblet cells in villi (Fig. 3J, K and Fig. EV3E-G) due to intestinal epithelial JWA deletion. Moreover, the mRNA levels of mature IECs markers further revealed the disordered composition of intestinal cell types caused by intestinal epithelial JWA deletion. Such as lower absorptive enterocyte markers (Sis, Lct, Dpp4) in villi (Fig. 3L and Fig. EV3H) and Paneth cell markers (Lyz1, Mmp7) in crypts (Fig. 3M and Fig. EV3H); higher goblet cell markers (Muc2, Spink4, Tuff3), enteroendocrine cell markers (Gcg, Ngn3, Reg4), and tuft cell markers (Dclk1, Hck) in villi (Fig. 3N and Fig. EV3H). Our results suggested intestinal epithelial JWA deletion might lead to secretory intestinal epithelial cell differentiation lineages, except Paneth cells.

### Intestinal epithelial JWA deletion exacerbates the injury and prevents intestinal epithelial regeneration after whole abdomen X-ray irradiation exposure

To investigate the regeneration capability of intestinal epithelium, we subjected the mice to whole abdomen X-ray irradiation (WAI) at a dose of 10 Gy. We sacrificed the mice at the indicated time points after WAI and collected crypts or paraffin sections from jejunum for later analysis (Fig. 4A and Fig. EV4A). We saw the rising of JWA in both mRNA and protein levels during the recovery period within five days after WAI, and returned to base levels on the seventh day (Fig. EV4B and C). In addition, interestingly, the mRNA levels of ISCs markers olfm4 and lgr5 altered consistently with JWA (Fig. EV4C). These further prompted us that JWA participated in the regeneration of ISC and intestinal epithelium. However, as we expected, intestinal epithelial JWA deletion exacerbated the injury and prevented the regeneration of intestinal epithelium. The body weights of JWA^IEC-ko^ mice lost faster than JWA^IEC-wt^ mice (Fig. 4B). We dissected the mice and found that JWA^IEC-ko^ mice showed visible intestinal hyperemia and swelling and markedly shorter intestine rather than JWA^IEC-wt^ mice (Fig. 4C and D). We performed the intestinal epithelial permeability assay in mice after WAI and found that the intestinal epithelial of JWA^IEC-ko^ mice were more permeable to FD4 than JWA^IEC-wt^ mice (Fig. 4E). We also observed worse damaged crypts and villi (Fig. 4F-H) and more cleaved caspase-3 (CC3) positive cells (Fig. 4I) in intestine sections from JWA^IEC-ko^ mice, revealing more severe intestinal epithelial injury than JWA^IEC-wt^ mice. Moreover, we found fewer ki67 positive regenerative crypts (Fig. 4J and K) and fewer olfm4 positive cells (Fig. 4L and M) in regenerative crypts owing to intestinal epithelial JWA deletion. These results suggested intestinal epithelial JWA deletion exacerbated the injury and prevented intestinal epithelial regeneration after WAI.

**Figure 4.**
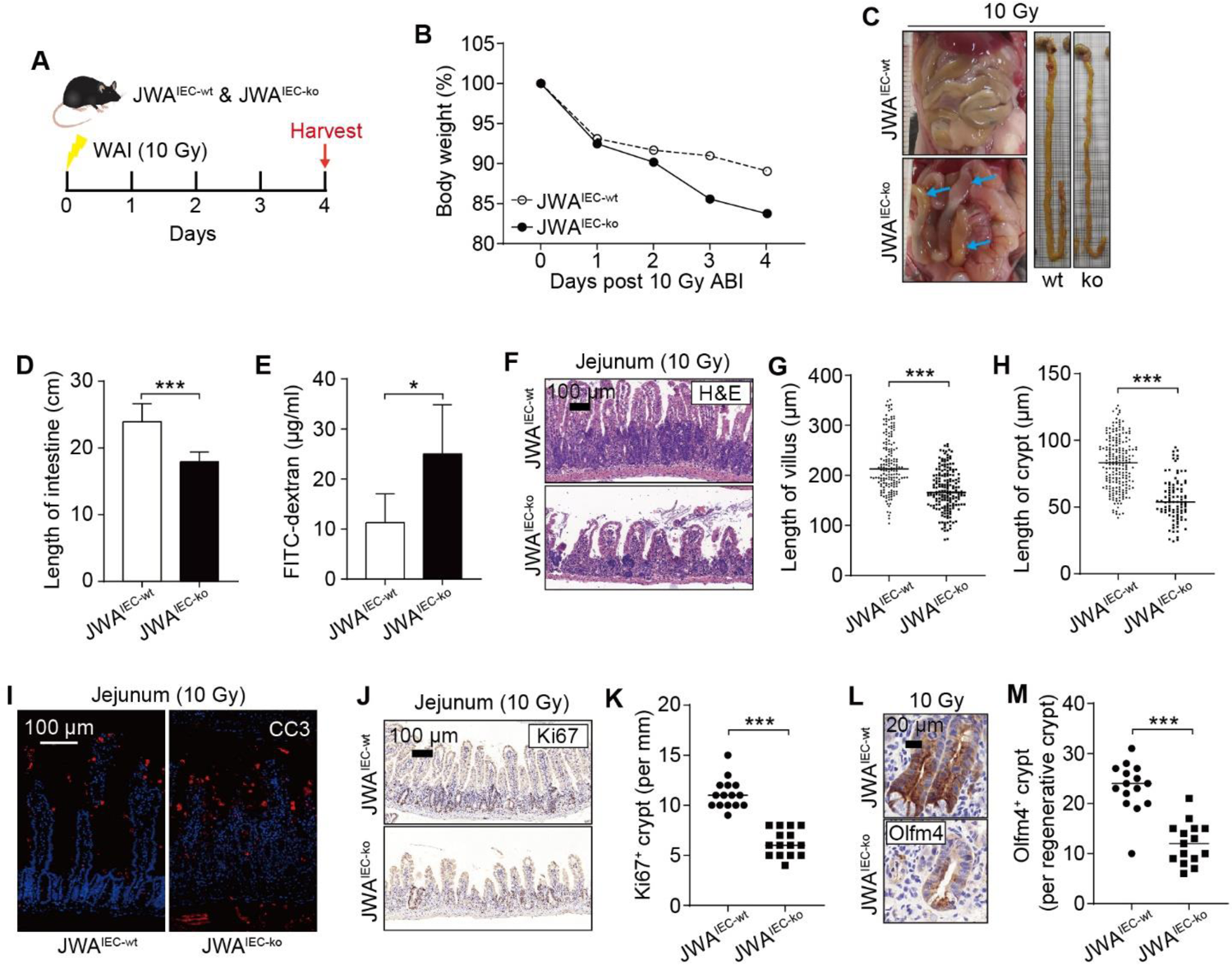
Intestinal epithelial JWA deletion exacerbates injury and prevents regeneration of intestinal epithelium after X-ray exposure. A. Timeline of the X-ray exposure model performed on JWA^IEC-wt^ and JWA^IEC-ko^ mice. B. Bodyweight curve of JWA^IEC-wt^ and JWA^IEC-ko^ mice after X-ray exposure; n=6 for JWA^IEC-wt^ mice and n=5 for JWA^IEC-ko^ mice. C. Representative abdominal cavity and intestinal images of JWA^IEC-wt^ and JWA^IEC-ko^ mice after X-ray exposure. D. Intestinal length of JWA^IEC-wt^ and JWA^IEC-ko^ mice after X-ray exposure; n=6 for each genotype. E. Evaluate of intestinal epithelial permeability to FD4 in the JWA^IEC-wt^ and JWA^IEC-ko^ mice after X-ray exposure; n=6 for each genotype. F-H. H&E staining of the jejunum sections of JWA^IEC-wt^ and JWA^IEC-ko^ mice after X-ray exposure (F), and the length measurement of villi (G) and crypts (H); n=3 for each genotype; scale bar: 100μm. I. Immunofluorescence staining of Cleaved caspase-3 in the intestinal sections of JWA^IEC-wt^ and JWA^IEC-ko^ mice after X-ray exposure; scale bar: 100 μm. J, K. Immunochemistry staining of ki67 in the intestinal sections of JWA^IEC-wt^ and JWA^IEC-ko^ mice after X-ray exposure (J), and the ki67 positive crypts counts (K); n=3 for each genotype; scale bar: 100 μm. L, M. Immunochemistry staining of olfm4 in the intestinal sections of JWA^IEC-wt^ and JWA^IEC-ko^ mice after X-ray exposure (L), and the olfm4 positive cells counts in the regenerative crypts (M); n=3 for each genotype; scale bar: 20 μm. Data information: In (B), data are presented as mean; In (D, E, G, H, K, M), data are presented as mean ± SD, \**P* <0.05 and \*\*\**P* <0.001 (Student’s *t*-test).

### JWA regulates PPARγ/Stat5 axis through Notch signal pathway

To explore the JWA deletion-induced molecule alterations, we distinguished the differential proteins in jejunum tissues between JWA^wt^ and JWA^ko^ mice through proteomics analysis (Dataset. EV1). We discovered that JWA deletion distinctively down-regulated the protein levels of signal transducer and activator of transcription 5 (Stat5) (Fig. 5A), which was intimately related to the maintenance of ISC and intestinal epithelium. We then determined the reduced levels of stat5 in both jejunum from JWA^ko^ mice and crypts from JWA^IEC-ko^ mice (Fig. 5B and C), nevertheless, we could neither detect the phosphorylated stat5 in jejunum nor in crypts. Therefore, we examined the upstream molecule of stat5, Janus Kinase 2 (JAK2), and phosphorylated JAK2 in crypts (Fig. 5C), which could phosphorylate stat5 at tyrosine residues. However, JWA deletion did not alter the phosphorylated levels of JAK2 in crypts (Fig. 5C), reminding that JWA deficiency might not affect the phosphorylation of stat5. According to the report, stat5 could face the transcriptional repression resulting from over-expression of the transcription factor peroxisome proliferator-activated receptor-gamma (PPARγ), which negatively affected ISCs maintenance. Results showed that JWA deletion did elevate the protein level of PPARγ and reduced the mRNA levels of stat5a and stat5b in crypts (Fig. 5C and D). Interestingly, JWA deletion also increased the mRNA level of PPARγ (Fig. 5D). We searched the JASPAR database (https://jaspar.genereg.net/) for the potential transcript factors (TFs) binding to the promoter region of PPARγ. We found that the primary TFs were the hairy and enhancer of splits (Hes), a target genes family of Notch signal (Appendix Table. S1). We quantified several Hes genes and found that JWA deletion drastically reduced the mRNA levels of hes1 (Fig. 5E), a transcription repressor of PPARγ; these results informed us that JWA deletion might cause Notch signal inhibition. We then confirmed that JWA deletion down-regulated the levels of the cleaved (Cl) Notch1, Notch1 intracellular domain (NICD), and Hes1 in crypts (Fig. 5F). However, the full-length (Fl) Notch1 showed no alterations in both protein and mRNA levels (Fig. 5F and G). Furthermore, we knocked down JWA levels in the rat intestinal epithelial cell line (IEC-6) by transfecting with short hairpin RNA of JWA (shJWA), and we observed the similar molecule changes as JWA deletion crypts (Fig. 5H). Fortunately, Hes1 over-expression and the PPARγ antagonist GW9662 could retrieve the changes (Fig. 5I, J). Our results suggested that JWA deficiency disturbed PPARγ/Stat5a axis by inhibiting the Notch signal.

**Figure 5.**
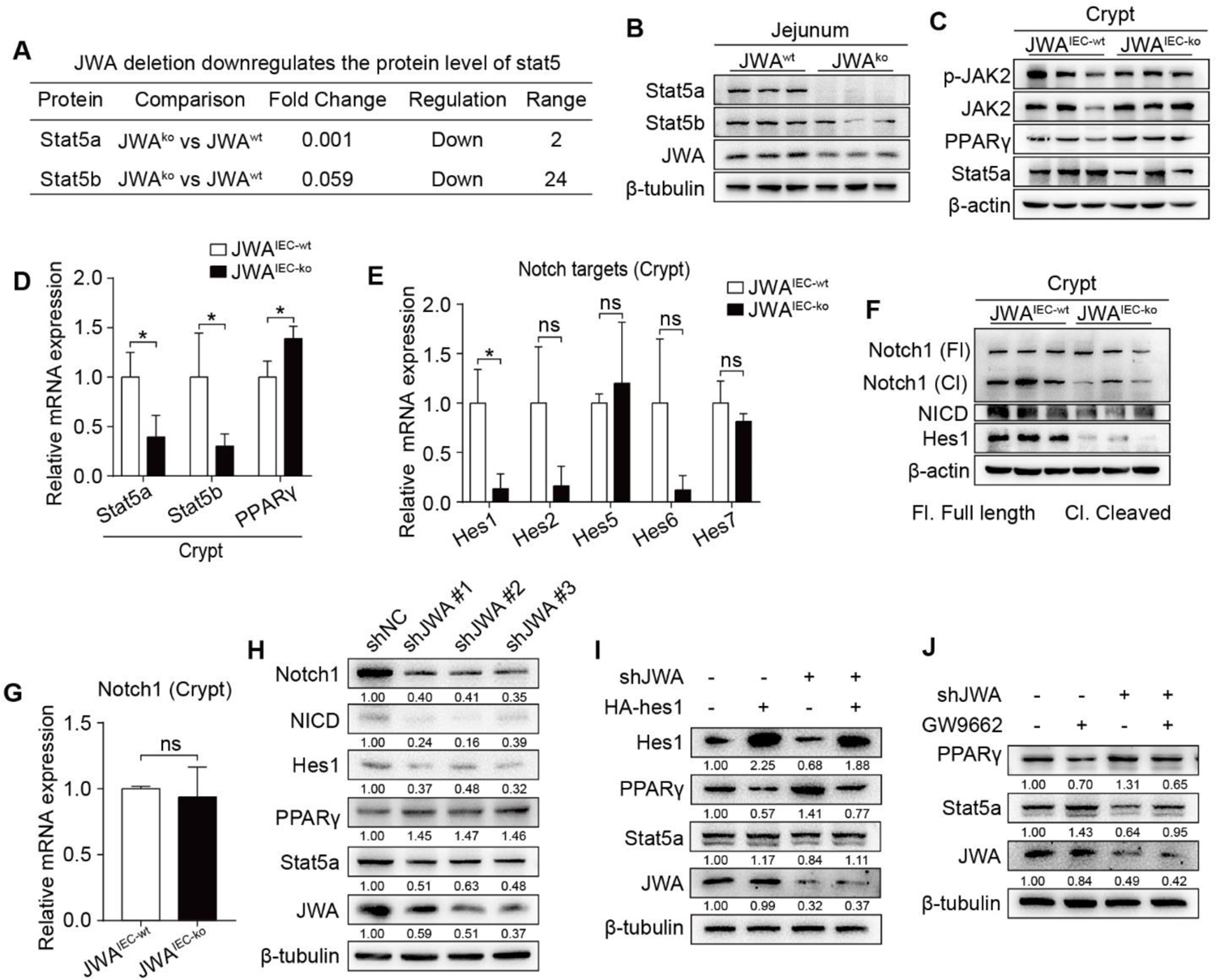
JWA regulates PPARγ/Stat5 axis through Notch signal pathway. A. Screen of the down-regulated molecules Stat5 from the proteomics analysis performed on the jejunum of JWA^wt^ and JWA^ko^ mice. B. Immunoblotting of Stat5a, Stat5b and JWA in the jejunum of JWA^wt^ and JWA^ko^ mice; n=3 for each genotype. C. Immunoblotting of PPARγ, Stat5a, Stat5b, JAK2 and p-JAK2 in the crypts of JWA^IEC-wt^ and JWA^IEC-ko^ mice; n=3 for each genotype. D. QRT–PCR detection of Stat5a, Stat5b and PPARγ levels in the crypts of JWA^IEC-wt^ and JWA^IEC-ko^ mice; n=3 for each genotype. E. QRT–PCR detection of Notch target genes hes1, hes2, hes5, hes6 and hes7 in the crypts of JWA^IEC-wt^ and JWA^IEC-ko^ mice; n=3 for each genotype. F. Immunoblotting of full-length Notch1, cleaved Notch1, NICD and hes1 in the crypts of JWA^IEC-wt^ and JWA^IEC-ko^ mice; n=3 for each genotype. G. QRT–PCR detection of Notch1 levels in the crypts of JWA^IEC-wt^ and JWA^IEC-ko^ mice; n=3 for each genotype. H. Immunoblotting of JWA, Stat5a, PPARγ, hes1, NICD and Notch1 in IEC-6 cells transfected with shJWA plasmids. I. Immunoblotting of JWA, Stat5a, PPARγ and hes1 in IEC-6 cells co-transfected with shJWA and HA-hes1 plasmids. J. Immunoblotting of JWA, Stat5a and PPARγ in IEC-6 cells transfected with shJWA plasmid followed by GW9662 (10 μM) treatment. Data information: In (D, E, G), data are presented as mean ± SD, \**P* <0.05 and \*\*\**P* <0.001 (Student’s *t*-test).

### JWA deficiency promotes ubiquitination degradation of Notch1 via ERK/Fbxw7 signal axis

To verify if JWA deficiency affected the stability of Notch1 protein, we conducted the cycloheximide (CHX)-chase assay in IEC-6 cells. Results showed that JWA deficiency accelerated Notch1 protein degradation (Fig. 6A and B). We then performed the *in vitro* ubiquitination assay and found that JWA deficiency promoted the degradation of Notch1 via ubiquitin-proteasome pathway (Fig. 6C). To gain insight into the reason for JWA deficiency-induced Notch1 degradation, we predicted the potential E3 ubiquitin ligases acting on Notch1 through the UbiBrowser Database (https://ubibrowser.ncpsb.org.cn/ubibrowser/). We screened the top five high-confidence scored ligases (Fig. EV5A) and verified them in JWA deficiency crypts and cells. However, JWA deficiency did not increase their levels in crypts or cells (Fig. EV5B and C). We have previously confirmed JWA as an upstream activator of the ERK/MAPK signal. Reportedly, ERK negatively regulates F-box and WD repeat domain containing-7 (Fbxw7), a known E3 ubiquitin ligase targets multiple substrates including Notch1. Fbxw7 is also a candidate in the predicted list although with low-confidence score. Therefore, we suspected that JWA deficiency might promote degradation of Notch1 through ERK/Fbxw7 axis. It was worth noting that we did observe that JWA deficiency inhibited phosphorylation of ERK1/2 and increased the levels of Fbxw7 in both crypts and cells (Fig. 6D and Fig. EV5D). To further clarify the role of the ERK/Fbxw7 signal axis on JWA deficiency-induced Notch1 degradation, we administrated the cells with Pamoic Acid (PA), an ERK agonist. The results showed that of ERK activation could effectively inhibit Fbxw7 expression, reverse JWA deficiency-induced Notch1 downregulation and downstream molecular changes (Fig. 6E and Fig. EV5E). Meanwhile, we transfected the cells with the small interfering RNA of Fbxw7 (siFbxw7) and achieved the same consequent as PA treatment (Fig. 6F and Fig. EV5F). Furthermore, we performed the CHX-chase assay and the *in vitro* ubiquitination assay in PA treated or siFbxw7 transfected cells. Results showed that PA treatment or siFbxw7 transfection could effectively enhance the stability of Notch1 and reduce the ubiquitination degradation of Notch1 caused by JWA deficiency (Fig. 6G-L). Our results suggested that JWA negatively regulates the E3 ubiquitin ligase Fbxw7 via ERK/MAPK signal and maintained the stability of Notch1. JWA deficiency promoted Notch1 degradation through the ubiquitin-proteasome pathway.

**Figure 6.**
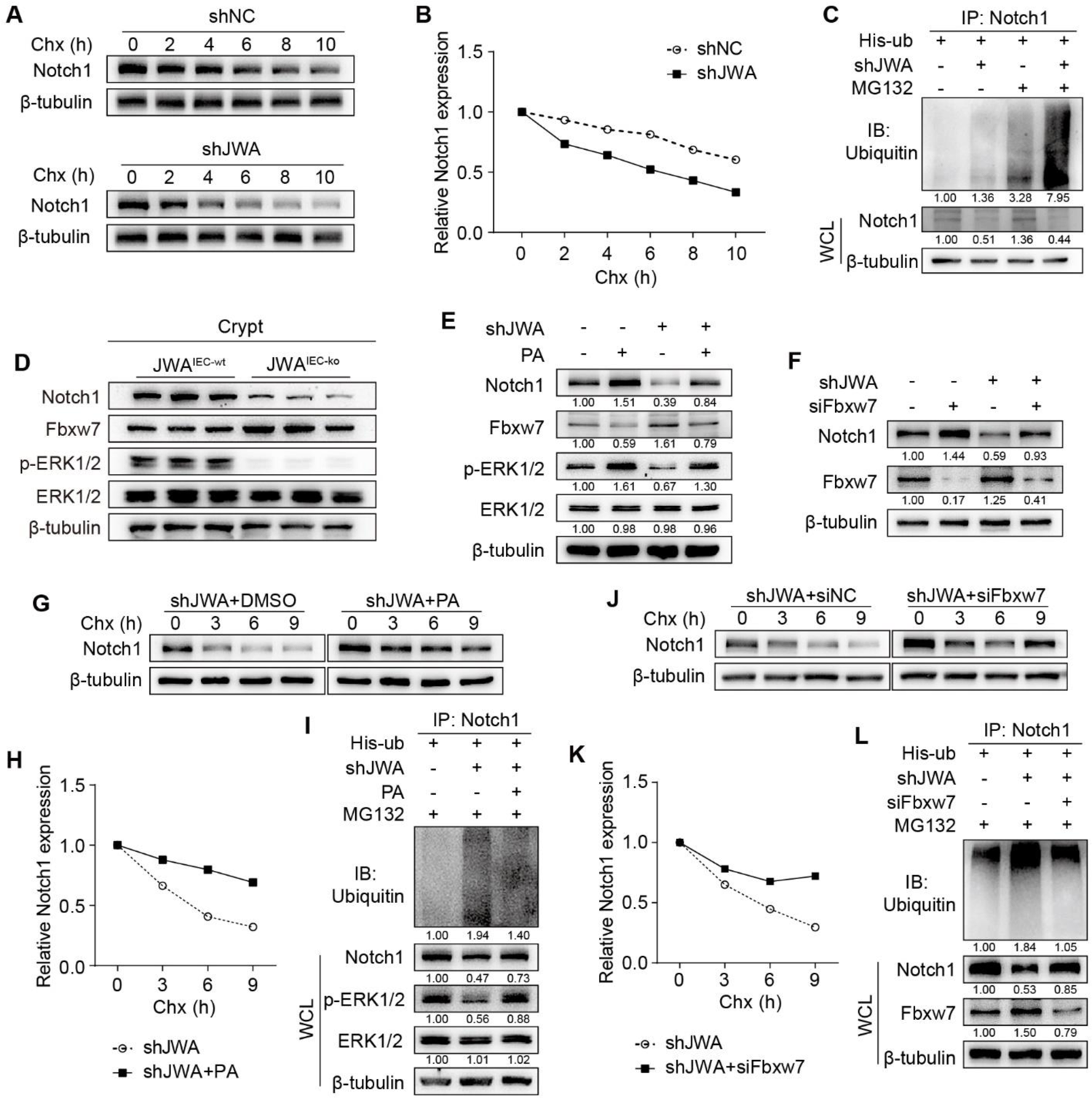
JWA deficiency promotes ubiquitination degradation of Notch1 through ERK/MAPK/Fbxw7 axis. A, B. CHX-chase assay in IEC-6 cells transfected with shNC or shJWA plasmid. C. *In vitro* ubiquitination assay of Notch1 protein in IEC-6 cells co-transfected with shJWA plasmid. D. Immunoblotting of Notch1, Fbxw7, ERK and p-ERK1/2 in the crypts of JWA^IEC-wt^ and JWA^IEC-ko^ mice; n=3 for each genotype. E. Immunoblotting of Notch1, Fbxw7, ERK and p-ERK1/2 in IEC-6 cells transfected with shJWA plasmids followed with Pamoic Acid (10 μM) treatment. F. Immunoblotting of Notch1 and Fbxw7 in IEC-6 cells co-transfected with shJWA plasmid and siFbxw7. G, H. CHX-chase assay in IEC-6 cells transfected with shJWA plasmid followed by Pamoic Acid (10 μM) treatment. I. *In vitro* ubiquitination assay of Notch1 protein in IEC-6 cells co-transfected with shJWA followed by Pamoic Acid (10 μM) treatment. J, K. CHX-chase assay in IEC-6 cells co-transfected with shJWA plasmid and siFbxw7. L. *In vitro* ubiquitination assays of Notch1 protein in IEC-6 cells co-transfected with shJWA and siFbxw7.

## Discussion

Aging is a progressive physiological process accompanying the age-dependent decline in physiological functions until the end of life. Aging is the result of gene and environment interactions. Excessive ROS produced by environmental stimuli can induce oxidative stresses and DNA damages; the body can resolve these issues through antioxidation and DNA repair pathways (Chesnokova *et al*, 2021; Lopez-Otin *et al*., 2013). However, deficiencies or mutants of the antioxidation and DNA repair genes such as NF-E2-related factor-2 (Nrf2) and X-Ray Repair Cross Complementing 1 (XRCC1) allow the accumulation of unrepaired DNA damage, inducing genomic instability and cell dysfunction or apoptosis, thus promoting aging (Wang *et al*, 2021; Zhao *et al*, 2021). Otherwise, increasing DNA repair capability facilitates longevity (Chesnokova *et al*., 2021). JWA is an environmental response gene upregulating under oxidative stress stimuli and regulates XRCC1 to repair the DNA single-strand breaks (Chen *et al*., 2007b; Wang *et al*., 2009). JWA also resists paraquat-induced oxidative stress in neurons by upregulating the Nrf2 level in Parkinson’s disease (PD) mice model (Zhao *et al*., 2017). The present study found that genetic JWA deletion induced multiple aging-related phenotypes and dramatically shortened lifespan in mice, suggesting that JWA was a new aging-associated gene.

Intestinal epithelium is a rapid renewal tissue composed of villi and crypts. The proliferative intestinal stem cells (ISCs) that reside in the bottom of crypts are responsible for epithelial maintenance. ISCs generate the transit-amplifying progenitor cells that differentiate into mature intestinal epithelial cells (IECs) and migrate up to the villi to replace senescent or damaged IECs and complete epithelial renewal (Stine *et al*, 2019). These characteristics make the intestinal epithelium more susceptible to environmental factors such as ionizing radiation (Chaves-Perez *et al*., 2019). Since the reduced ISCs activity, the intestinal epithelium regenerates more slowly in old mice than young mice after irradiation-induced injury (He *et al*, 2020; Pentinmikko *et al*., 2019). The present study found intestinal epithelial JWA deletion dramatically inhibited the proliferating of crypt cells and reduced the number of lgr5^+^ ISCs, thus causing epithelial atrophy, which was more noticeable in elderly mice. Moreover, we found that JWA responded to X-ray irradiation and participated in epithelial regeneration. Intestinal JWA deletion aggravated injury and prevented regeneration of the epithelium in mice after 10 Gy X-ray exposure; these phenotypes were similar to aging mice. Our results suggested that JWA was a regulator for intestinal epithelial maintenance and aging.

An important question is why JWA deficiency accelerates aging, considering the long-term nutrient malabsorption through depauperated intestinal epithelium. Dietary restriction (DR) without malnutrition is beneficial for improving aging and aging-related diseases. For example, reducing protein or specific EAAs is associated with longevity (Wang *et al*., 2022). However, nutrition like EAAs is a double-edged sword in health maintenance. Restriction and supplementation of specific EAAs both show promising outcomes in expanding lifespan. Nonetheless, long-term restriction or deficiency of EEAs may lead to adverse events, such as growth retardation and aging acceleration (Navik *et al*, 2021). In this study, JWA gene deletion resulted in intestinal epithelial dysplasia from the embryonic stage to the whole life cycle of mice. Intestinal epithelium is the predominant area for nutrient absorption. Therefore, epithelial atrophy may undoubtedly cause long-term malnutrition, thus inducing stunted growth and promoting aging. The present study showed that JWA deletion reduced food, water intake and intestinal absorption, caused light weight and thin body in mice. These results were parallel to the existing report on mice lacking the Glutamate transporter associated protein 3-18 (GTRAP3-18), an alternate name of JWA (Aoyama *et al*, 2018). Although JWA deletion-induced intestinal maldevelopment in mice embryo, it did not alter the embryo weight due to the independent of intestinal absorption in embryonic growth. Our results suggested that JWA deficiency-induced intestinal epithelial atrophy might cause chronic malnutrition and accelerated aging. To detailly explain the reason for JWA deficiency accelerated aging, more research programs, such as epithelial absorption and metabonomic analysis for nutrients, need to conduct.

Multiple signaling transduction pathways, including Wnt, Notch, EGF, BMI, YAP/TAZ, etc., independently or cooperatively maintain intestinal epithelial homeostasis through ISC dependent renewal and regeneration (Beumer & Clevers, 2021). JAK/STAT signaling pathway is a central node that regulates cellular processes through signal transduction (Hu *et al*, 2021). It responds to tissue turnover and regeneration via regulating stem cells, including hematopoietic stem cells (Holdreith *et al*, 2021), embryonic stem cells (Lin *et al*, 2021), as well as ISCs (Du *et al*, 2020). It is also a maintainer for cancer stem cells and promoter for multiple cancers, such as colorectal cancer (Silva *et al*, 2021). Stat5, consisting of the transcription activator Stat5a and Stat5b, is a positive contributor to ISC and intestinal epithelial regeneration in the dextran sulfate sodium (DSS)-induced colitis, *Clostridium difficile* infection-induced Ileocolitis, and irradiation-induced intestinal injury mouse models (Gilbert *et al*, 2015; Liu *et al*, 2019). In the present study, our preliminary discovery on the mechanism was that JWA deficiency down-regulated Stat5 at both the transcription and translation levels, suggesting that JWA maintains intestinal epithelium through Stat5 signaling.

PPARs, consist of three subtypes (α, δ/β, and γ), are nuclear hormone receptors activated by fatty acids and endogenous ligands, and responsible for lipid and glucose metabolism homeostasis (Mihaylova *et al*, 2018; Trauner & Fuchs, 2022). PPARα and PPARδ/β is able to augment ISC function in high-fat diet, aging and injury (Mana *et al*, 2021; Mihaylova *et al*., 2018), and promotes intestinal tumorigenesis (Mana *et al*., 2021). PPARγ plays bidirectional roles in ISC and epithelial maintenance. On the one hand, PPARγ regulates the proliferation and differentiation of intestinal progenitor cells through the region-specific promotion of fatty acid oxidation, and maintains intestinal epithelial renewal (Stine *et al*., 2019). On the other hand, high PPARγ activity shows the inhibitory effect on Wnt/β-catenin signal pathway and impairs Lgr5^+^ ISC function (Pereira *et al*, 2020), and leads to colon cancer stem cells inhibition (Moon *et al*, 2014). Moreover, PPARγ activation inhibits various signals, including JAK/STAT (Vallee *et al*, 2018). The inhibitory effect on leukemia stem cells works by suppressing Stat5 expression (Kumar *et al*, 2020). The present study showed that JWA deficiency elevated both transcription and translation levels of PPARγ, considering it a reason for Stat5 reduction.

The Notch signal pathway regulates cell proliferation, differentiation, tumorigenesis, and stem cell maintenance (Siebel & Lendahl, 2017; Silva *et al*., 2021). Transcription repressor Hes1 is a target gene of Notch signal and plays an essential role in stem and progenitor cell maintenance; it is reported to prevent hematopoietic stem cell exhaustion by suppressing PPARγ expression (Ma *et al*, 2020; Wu *et al*, 2021). Moreover, Notch/Hes1 axis inhibition accompanied by PPARG activation can repress the progression of colorectal cancer via suppressing cancer stem cells (Moon *et al*., 2014). Here we reported that JWA deficiency inhibited Notch signal through ubiquitin-proteasome pathway-mediated degradation of the Notch1 receptor. The down-regulation of Notch target gene Hes1 was also the reason for PPARγ activation due to JWA deficiency. The E3-ubiquitin ligase Fbxw7 targets multiple substrates such as c-jun, c-myc, and Notch1; it works as a tumor suppressor for colorectal cancer by inhibiting Notch signal. However, Fbxw7 deletion alters the maintenance of intestinal stem/progenitor cells and the fate of differentiated cells and induces colorectal tumorigenesis via losing suppression on Notch in mice intestine (Babaei-Jadidi *et al*, 2011; Sancho *et al*, 2010). The tumor suppressor Fbxw7 expression is inversely correlated with ERK activation in pancreatic cancer, sustained activation of the Ras-Raf-MEK-ERK pathway phosphorylates Fbxw7 at Thr205, and destabilizes Fbxw7 (Ji *et al*, 2015; Ye *et al*, 2021). We previously demonstrated that JWA was an upstream activator of the ERK/MAPK signal pathway (Chen *et al*, 2007a; Mao *et al*, 2006). The present study showed that JWA deficiency up-regulated Fbxw7 expression by inhibiting the phosphorylation of ERK1/2, thus promoting Notch1 degradation via ubiquitin-proteasome pathway. However, more exact mechanism by which JWA regulates the Notch signal pathway need further investigation.

Notch signal promotes the IECs differentiation into enterocyte lineage while antagonizing secretory cell fate in intestine (Ludikhuize *et al*, 2020; Sancho *et al*., 2010), which is reversed by Fbxw7 deletion (Sancho *et al*., 2010). The present study found that intestinal JWA deletion promoted differentiated IECs into secretory cell lineages and suppressed the absorbing cell lineages. However, Paneth cells were reduced, presumably due to the suppression of Paneth cells by JWA deficiency-caused stat5 down-regulation (Liu *et al*., 2019). Therefore, the typical phenotype of Notch signal inhibition, co-localization of Paneth cell marker Lysozyme and Goblet cell marker Muc2 in crypts (Stine *et al*., 2019), was not observed in JWA^IEC-ko^ mice. The reason for JWA deficiency-caused differential regulation on Paneth cells and other secretory cell lineages is an interesting issue worthy of further exploration.

In summary, the present study verified that JWA deficiency accelerated aging through disrupting intestinal epithelial homeostasis. Mechanistically, we found that JWA deficiency promoted Notch1 degradation via the ubiquitin-proteasome pathway in the ERK/Fbxw7 dependent manner, thus disturbing the PPARγ/Stat5 axis. This is the reason for ISC function decline and IEC lineages skew. It will now be important to expound how JWA deficiency-induced intestinal epithelial atrophy accelerating aging, whether it is due to nutrient malabsorption or other complex issues. Our results may also provide the intestinal epithelial homeostasis maintenance as a potential anti-aging strategy. Moreover, JWA may also be a potential intervention target for stem cell-driven colorectal cancer, although JWA is a suppressor for gastric and breast cancer (Ren *et al*, 2021; Zhang *et al*, 2021).

## Materials and Methods

### Mouse

JWA^flox/flox^, JWA^wt^ and JWA^ko^ mice were described in our previous study (Gong *et al*, 2012). Villin-cre mouse on a C57BL/6 background was obtained from Shanghai Model Organisms Center (Shanghai, China). Wild type C57BL/6 mouse was purchased from SLAC Laboratory Animal Co., Ltd (Shanghai, China). JWA^IEC-ko^ mouse was generated by cross-mating Villin-cre mouse with JWA^flox/flox^ mouse. All mice were maintained in the Animal Core Facility of Nanjing Medical University in a specific pathogen-free (SPF) condition. The animal operations were approved by the Institutional Animal Care and Use Committee of Nanjing Medical University (Approval No. IACUC-2004044).

### SA-β-gal staining

Freshly isolated liver tissue was embedded in the Tissue-Tek O.C.T compound (Sakura, Tokyo, Japan) and quick-frozen on dry ice. The 10 μm thickness frozen section was prepared on the CM1950 cryostat (Leica Biosystem, Wetzlar, Germany) and stained using a Senescent cell β-galactosidase staining kit (Servicebio, Wuhan, China) according to the manufacturer’s guidelines.

### Intestinal absorption assay by oral glucose tolerance test

Mouse was orally administrated with 6 mg/kg glucose. The blood glucose levels at indicated time points were measured using a handheld blood glucometer (Sinocare, Changsha, China) through tail tips.

### BrdU^+^ cell migration assay

Mouse was intraperitoneally injected with 100 mg/kg BrdU (APExBIO, Houston, TX, USA) dissolved in saline. The jejunum was isolated and fixed in 4% paraformaldehyde 3 hours after injection. The BrdU^+^ cells were detected by immunofluorescence using the BrdU antibody.

### Serum oxidoreductase activity detection

The whole blood was collected from the post-ocular vein of mouse and centrifugated to separate the serum. Enzymatic activities were measured using the Total Superoxide Dismutase (SOD) Assay Kit (Beyotime, Shanghai, China), Catalase (CAT) Detection Kit, and Glutathione Peroxidase (GSH-Px) Detection Kit (Aifang Biological, Changsha, China) according to the manufacturer’s protocol.

### Intestinal villus and crypt isolation

Mouse intestine was isolated and flushed with phosphate-buffered saline (PBS) supplemented with penicillin/streptomycin. The jejunum was opened longitudinally and scraped by coverslip to separate the villus composition, which was washed several times in PBS with penicillin/streptomycin. The remained jejunum was washed several times in PBS and cut into 5 mm pellets, crypts were isolated by shaking the pellets violently in PBS supplemented with 5 mM EDTA.

### X-ray exposure

Mouse was anesthetized and placed into the RS 2000 Pro X-ray irradiator (Rad Source Technologies, Buford, GA, USA) to receive X-ray exposure (10 Gy, 1.5 Gy/min), the exposure area was limited to the whole abdomen by the control of built-in beam limiter in the instrument. Mouse was sacrificed on the indicated days after irradiation, the blood and intestine were collected for further experiment.

### Intestinal permeability assay

Mouse was orally administrated with 15 mg fluorescein isothiocyanate isomer-dextran 4 kDa (FD4, Sigma-Aldrich, St. Louis, MO, USA), serum was collected 4 hours later. The total fluorescence intensity (Excitation: 485 nm, Emission: 528 nm) in serum was detected on the Infinite M200 PRO fluorescence microplate reader (TECAN, Männedorf, Switzerland). The concentration of FD4 was calculated via a standard curve prepared with serial dilutions of FD4.

### Histology, immunohistochemistry, and immunofluorescence

Isolated tissue was fixed in 4% paraformaldehyde and embedded in paraffin. The 6 μm thickness section was prepared on the Microm HM 340 E rotary microtome (Thermo Scientific, Waltham, MA, USA), and dewaxed in xylene and serial concentrations of ethanol before staining. For hematoxylin-eosin (H&E) and Alcian blue staining, the dewaxed section was stained using the H&E staining kit or Alcian blue staining kit (Servicebio, Wuhan, China) according to the manufacturer’s protocol. For immunohistochemistry, the dewaxed section was boiled in Tris-EDTA antigen retrieval buffer (0.01 M Tris, 1 mM EDTA, 0.05% Tween 20, pH 9.0) for 10 min and incubated with the primary antibody (Appendix Table S2) at 4℃ overnight after being blocked with 5% bovine serum albumin (BSA). Then the section was incubated with horseradish peroxidase (HRP)-labeled secondary antibody and stained using the DAB Color Development Kit (Servicebio, Wuhan, China). The section was permanently mounted using the glycerol jelly mounting medium (Servicebio, Wuhan, China) after dehydrated and clarified in ethanol and xylene. For immunofluorescence, the section was incubated with Alexa Fluor dye-labeled secondary antibody and mounted using the ProLong Diamond Antifade Mountant with DAPI (Thermo Scientific, Waltham, MA, USA). The images were obtained using the Pannoramic MIDI digital slide scanner system (3DHISTECH, Budapest, Hungary).

### Proteomics analysis

Freshly isolated jejunum was flushed with PBS with penicillin/streptomycin, and quick-frozen in liquid nitrogen. The protein spectrum analysis was performed by CapitalBio Technology (Beijing, China).

### Cell culture, transfection, and treatment

Rat small intestinal epithelial crypt (IEC-6) cell (Zhong Qiao Xin Zhou, Shanghai, China) was cultured in Dulbecco’s modified eagle medium supplemented with 10% fetal bovine serum (TransGen Biotech, Beijing, China), 100U/ml penicillin, 0.1mg/ml streptomycin (Biyotime, Shanghai, China) and 10 μg/ml recombinant human insulin (Zhong Qiao Xin Zhou, Shanghai, China). For cell transfection, shRNA plasmids for JWA (Keris, Nanjing, China), siRNA for Fbxw7 (GenPharma, Shanghai, China) and their nonspecific control were synthesized (Appendix Table S3), and the commercial HA-hes1 plasmid was purchased (Youbio, Changsha, China). The plasmids and siRNA were transfected or co-transfected using the Lipo8000 transfection reagent (Biyotime, Shanghai, China) following the manufacturer’s protocol. To activate the phosphorylation of ERK1/2 or suppress the expression of PPARγ, the cell was treated with 10 μM pamoic acid (Macklin, Shanghai, China) or 10 μM GW9662 (MedChemExpress, Monmouth Junction, NJ, USA) for 24 hours. For the CHX-chase assay, the cell was treated with 100 μg/ml cycloheximide (CHX) at the indicated time points.

### Immunoblotting

Total protein was extracted using the RIPA lysis buffer (50mM Tris, 150mM NaCl, 1% Triton X-100, 1% sodium deoxycholate, 0.1% SDS, pH 7.4) supplemented with protease and phosphatase inhibitor cocktail (NCM, Suzhou, China) and quantified using the bicinchoninic acid assay (BCA) kit (Beyotime, Shanghai, China). Protein (20-40 μg/lane) was separated by sodium dodecyl sulfate-polyacrylamide gel electrophoresis (SDS-PAGE) and blotted onto polyvinylidene fluoride (PVDF) membrane. The membrane was probed with diluted primary antibody (Appendix Table S2) at 4 °C overnight after being blocked in 5% non-fat powder milk. The blot was detected using an Enhanced Chemiluminescence (ECL) detection kit (Vazyme, Nanjing, China) on the Chemiluminescence Image Analysis System (Tanon, Shanghai, China) after the HRP-labeled secondary antibody incubation.

### Quantitative reverse transcription and polymerase chain reaction (qRT-PCR)

Total mRNA was isolated using the RNAiso Plus reagent (Takara, Beijing, China) and reverse transcribed using the HiScript II 1st Strand cDNA Synthesis Kit with gDNA wiper (Vazyme, Nanjing, China). The PCR reaction procedure was carried out on the ABI 7900HT Real-Time PCR System (Applied Biosystems, Carlsbad, CA, USA) with the AceQ qPCR SYBR Green Master Mix (Vazyme, Nanjing, China) following the manufacturer’s instruction. The synthetic cDNA was used as the template in the reaction, the primer pairs were listed in Appendix Table S4.

### *In vitro* ubiquitination assay

The cell was transfected with ubiquitin plasmid and treated with 10 μM MG132 (Selleck, Houston, TX, USA) for 6 hours. Total protein was then isolated using RIPA lysis buffer. Approximately 500 μg/250 μl protein per sample was incubated with the Notch1 antibody at 4 °C overnight and followed by incubated with Protein A/G Plus-Agarose (Beyotime, Shanghai, China) at 4 °C overnight. After being washed in PBS and centrifuged, the ubiquitin level in the precipitation was detected by immunoblotting.

## Statistical analysis

Data was analyzed on GraphPad Prism 8.0 software with the two-tail independent samples student’s *t* test, log-rank test, or Fisher’s exact test according to demand. The number of sample (n) was indicated in the figure legends. Data was presented as mean with standard deviation (mean ± SD) in the figures. *P* <0.05 was defined as the statistical difference.

## Data availability

Source data supporting this study was available upon request.

## Acknowledgements

This study was supported by the National Natural Science of China (Grant number: 81973156, 81673219).

## Author contributions

JWZ and XLi designed the research; JWL, LMW, YZ and YFW maintained and genotyped the mice; KD, LZ and XLiu assisted with experiments; APL managed the laboratory and assisted JWZ and XLi; YW and HLF assisted with the phenotype analysis; MH and GXD provided advises in aging research field; XLi and JWZ drafted and revised the manuscript; JWZ was the leading principal investigator in this research project.

---

**Synopsis.**
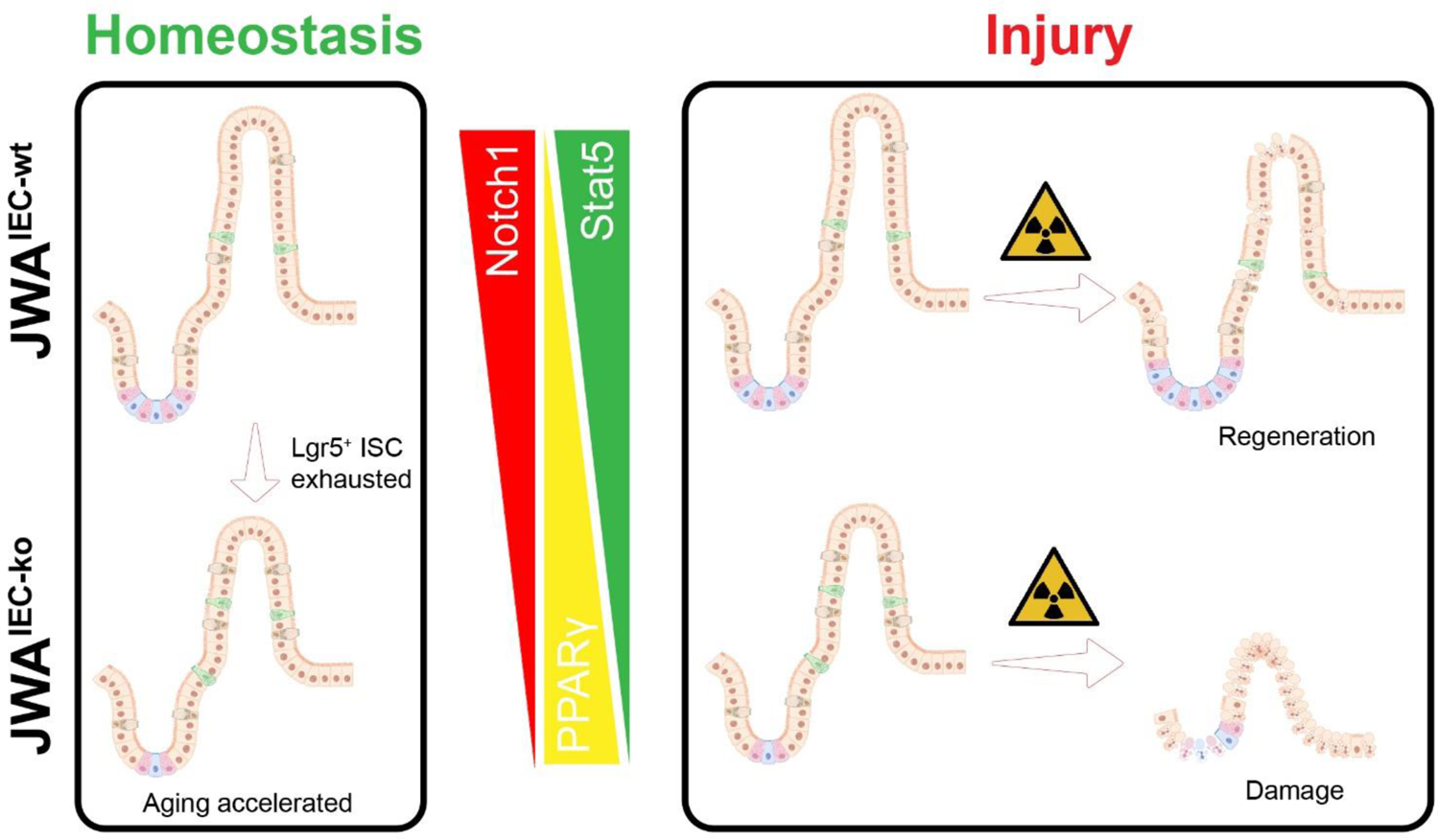

## Expanded View Figure Legends

**Figure EV1.**
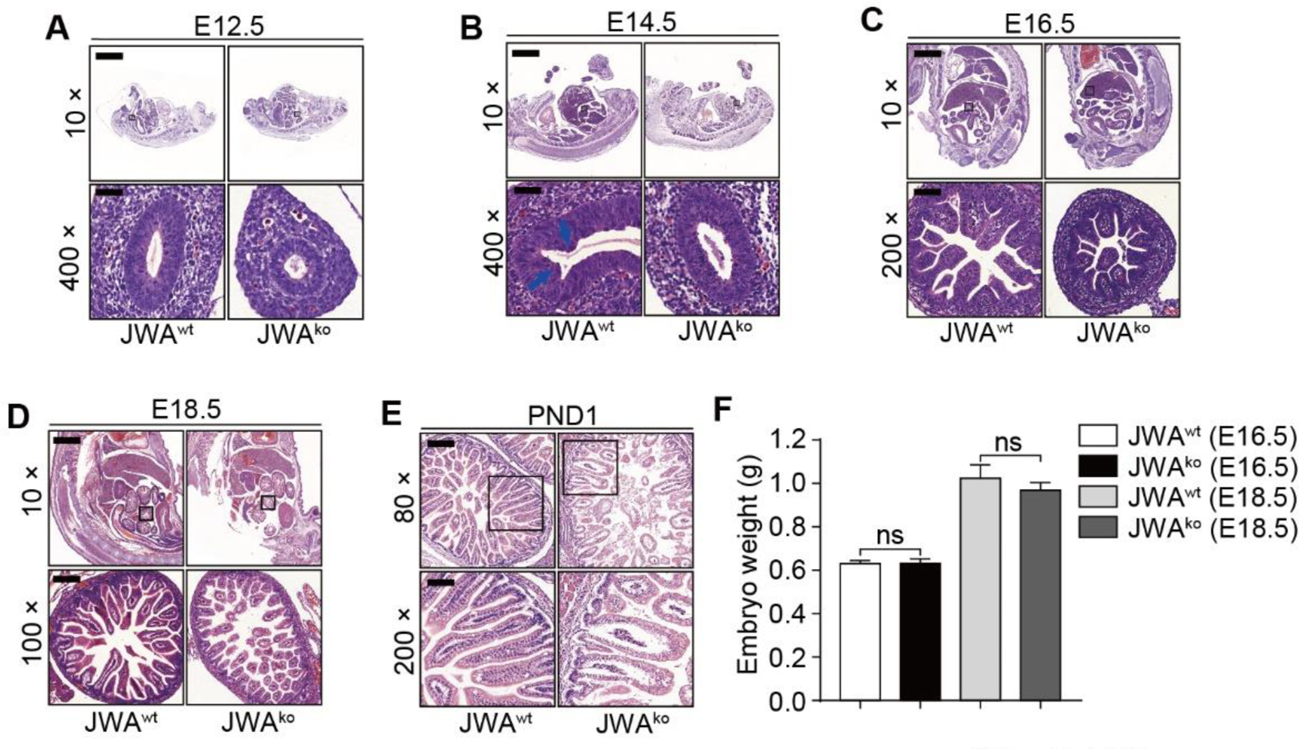
JWA deletion delays the early development of intestinal epithelium in mice A-D. H&E staining of the mice embryo sections at embryonic day E12.5 (A), E14.5 (B), E16.5 (C) and E18.5 (D); Scale bar: 1600 μm for 10×, 160 μm for 100×, 80 μm for 200× and 40 μm for 400×. E. H&E staining of the intestine in newborn mice on the postnatal day 1 (PND1); Scale bar: 200 μm for 80× and 80 μm for 200×. F. Embryonic weight of the mice embryo at E16.5 and E18.5. Data information: In (F), data are presented as mean ± SD, ^ns^ No significance (Student’s *t*-test).

**Figure EV2.**
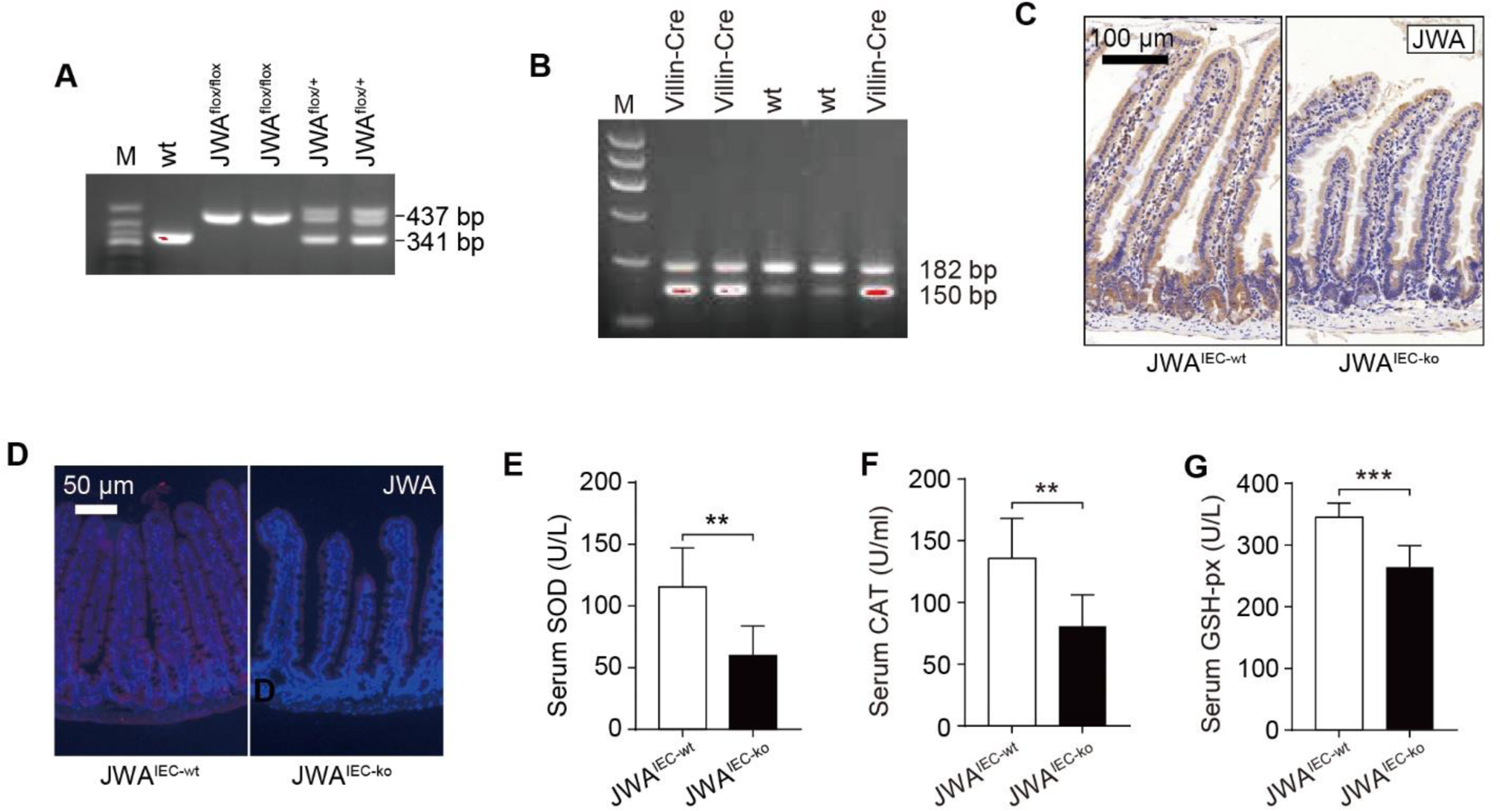
Intestinal epithelial JWA deletion reduces the antioxidant enzyme activity in aged mice. A, B. Agarose gel electrophoresis images of mouse genotyping. C. Immunochemistry staining of JWA in the intestinal sections of JWA^IEC-wt^ and JWA^IEC-ko^ mice; scale bar: 100 μm. D. Immunofluorescence staining of JWA in the intestinal sections of JWA^IEC-wt^ and JWA^IEC-ko^ mice; scale bar: 50 μm. E-G. Activities of the antioxidant enzymes SOD (E), CAT (F) and GSH-px (G) in the serum of 10-month-old JWA^IEC-wt^ and JWA^IEC-ko^ mice. Data information: In (E-G), data are presented as mean ± SD, \*\**P* <0.01 and \*\*\**P* <0.001 (Student’s *t*-test).

**Figure EV3.**
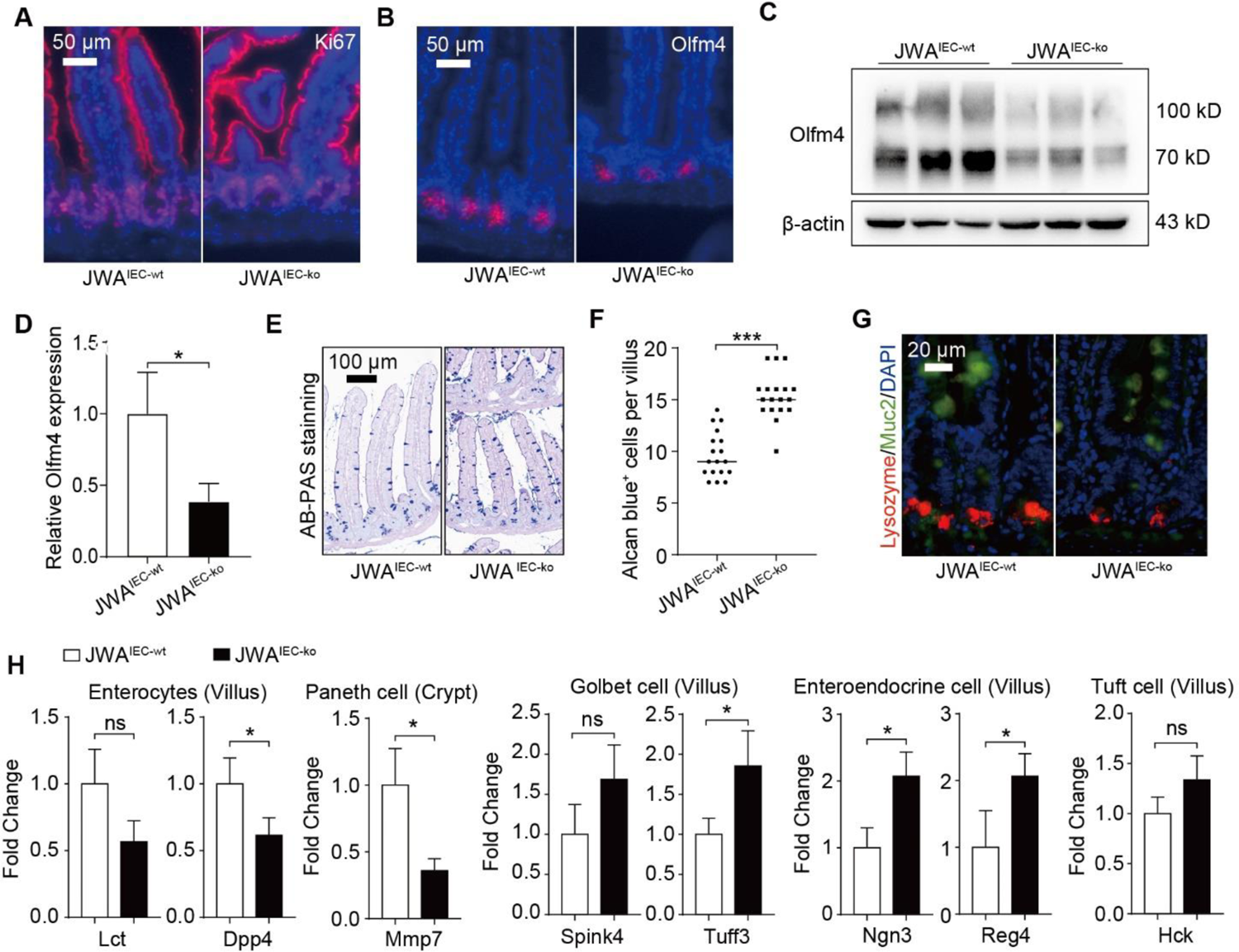
Intestinal epithelial JWA deletion reduces intestinal stem cells and skews the distributions of intestinal epithelial cells lineage. A. Immunofluorescence staining of ki67 in the intestinal sections of JWA^IEC-wt^ and JWA^IEC-ko^ mice; scale bar: 50 μm. B. Immunofluorescence staining of olfm4 in the intestinal sections of JWA^IEC-wt^ and JWA^IEC-ko^ mice; scale bar: 50 μm. C, D. Immunoblotting of olfm4 (C) and relative olfm4 levels (D) in the crypts of JWA^IEC-wt^ and JWA^IEC-ko^ mice; n=3 for each genotype. E, F. Alcian blue staining for the goblet cells (E) and counts for the AB positive cells (F) in villus of JWA^IEC-wt^ and JWA^IEC-ko^ mice; n=3 for each genotype; Scale bar: 100 μm. G. Immunofluorescence co-staining of lysozyme and muc2 in the intestinal sections of JWA^IEC-wt^ and JWA^IEC-ko^ mice; scale bar: 20 μm. H. QRT–PCR detection of the absorption enterocytes markers (Lct, Dpp4) in villi; Paneth cell marker (Mmp7) in crypts; goblet cell markers (Spink4, Tuff3), enteroendocrine cell markers (Ngn3, Reg4), and tuft cell marker (Hck) in villi; n=3 for each genotype. Data information: In (D, F, H), data are presented as mean ± SD, ^ns^ No significance, \**P* <0.05, \*\**P* <0.01 and \*\*\**P* <0.001 (Student’s *t*-test).

**Figure EV4.**
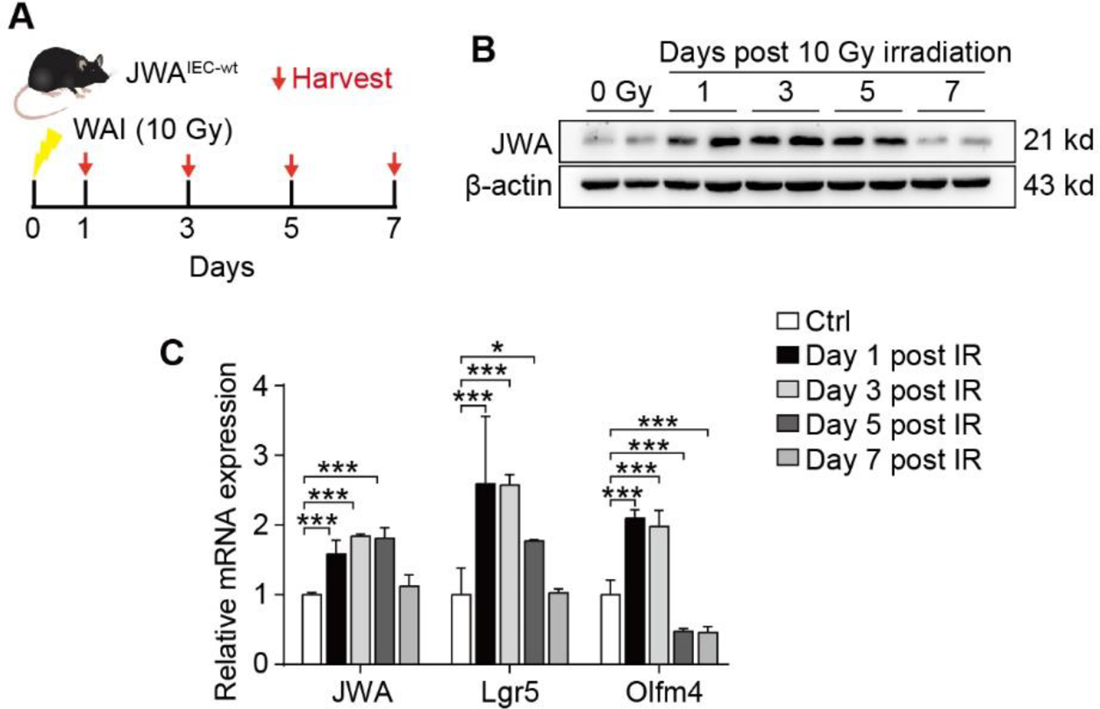
JWA participates in intestinal stem cell regeneration after X-ray exposure. A. Timeline for the X-ray exposure model performed on wild-type mice. B. Immunoblotting of JWA in the crypts of the wild-type mice at the indicated days after X-ray exposure. C. QRT–PCR detection of JWA and the ISCs markers (lgr5, olfm4) levels in crypts of the wild-type mice at the indicated days after X-ray exposure; n=6 for each time point. Data information: In (C), data are presented as mean ± SD, ^ns^No significance, \**P* <0.05, \*\**P* <0.01 and \*\*\**P* <0.001 (Student’s *t*-test).

**Figure EV5.**
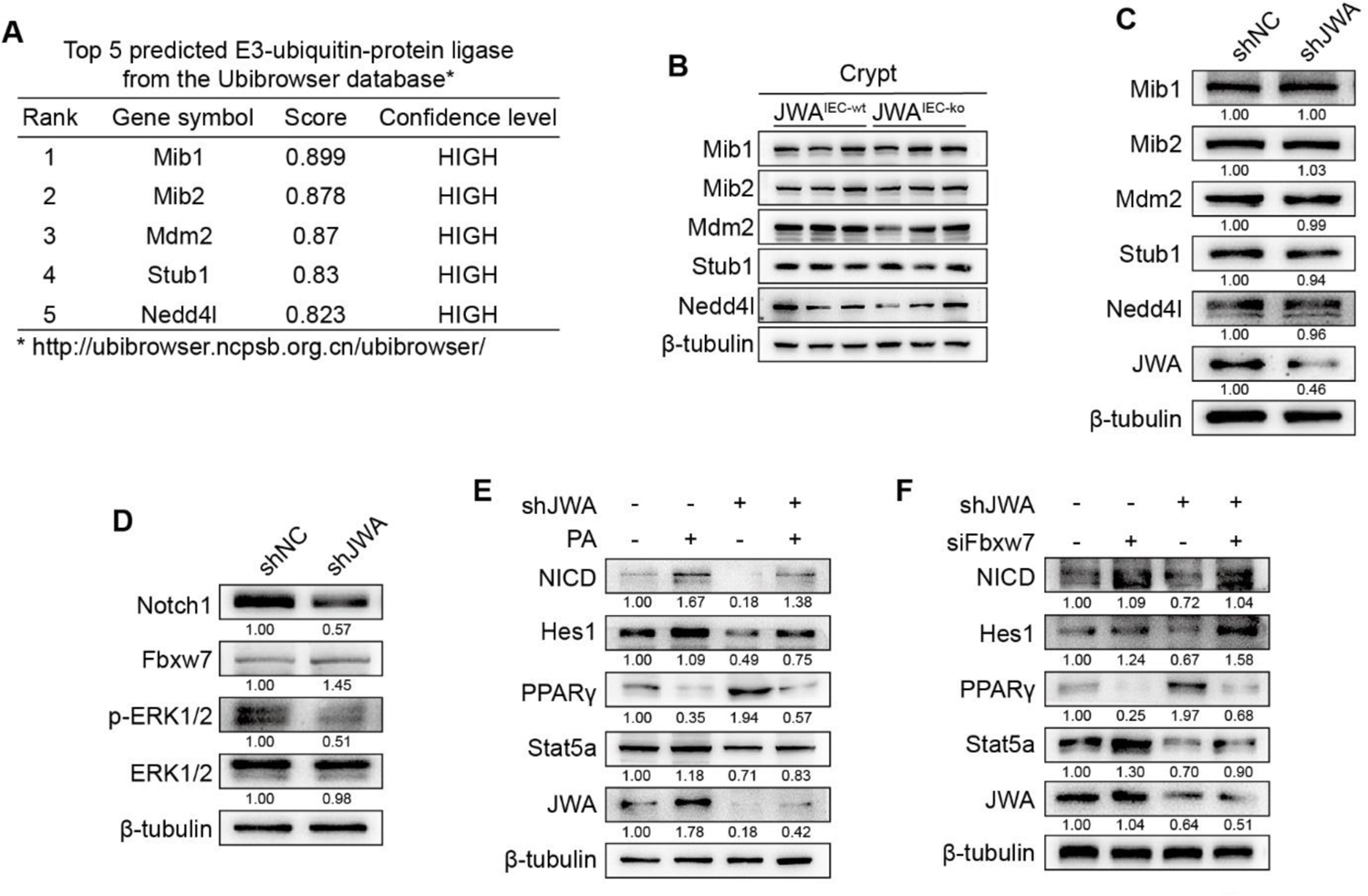
JWA deficiency promotes ubiquitination degradation of Notch1 through ERK /Fbxw7 and interferes with PPARγ/Stat5 axis. A. The predicted top five E3 ubiquitin ligases (Mib1, Mib2, Mdm2, Stub1 and Nedd4l) target Notch1. B. Immunoblotting of the predicted top five E3 ubiquitin ligases in crypts of JWA^IEC-wt^ and JWA^IEC-ko^ mice. C. Immunoblotting of the predicted top five E3 ubiquitin ligases in IEC-6 cells transfected with shJWA plasmid. D. Immunoblotting of Notch1, Fbxw7, ERK1/2 and p-ERK1/2 in IEC-6 cells transfected with shJWA plasmid. E. Immunoblotting of JWA, NICD, hes1, PPARγ and stat5a in IEC-6 cells transfected with shJWA plasmid followed by Pamoic Acid (10 μM) treatment. Immunoblotting of JWA, NICD, hes1, PPARγ and stat5a in IEC-6 cells co-transfected with shJWA plasmid and siFbxw7.

